# Prevalence of *RFC1*-Mediated Spinocerebellar Ataxia in a United States Ataxia Cohort

**DOI:** 10.1101/790006

**Authors:** Dona Aboud Syriani, Darice Wong, Claudio M. De Gusmao, Sameer Andani, Yuanming Mao, Giacomo Glotzer, Paul J. Lockhart, Sharon Hassin-Baer, Vikram Khurana, Soma Das, Christopher M. Gomez, Susan Perlman, Brent L. Fogel

## Abstract

**Objective:** Repeat expansions in *RFC1 and DAB1* have recently been identified as causing cerebellar ataxia, neuropathy, and vestibular areflexia syndrome (CANVAS) and spinocerebellar ataxia 37 (SCA37), respectively. We evaluated the prevalence of these repeat-expansions in an undiagnosed ataxia cohort from the United States.

**Methods:** A cohort of 596 patients with undiagnosed familial or sporadic cerebellar ataxia were evaluated at a tertiary referral ataxia center and excluded for common genetic causes of cerebellar ataxia. Patients were then screened for the presence of pathogenic repeat expansions in *RFC1* (AAGGG) and *DAB1* (ATTTC) using fluorescent repeat primed polymerase chain reaction (RP-PCR). Two additional undiagnosed ataxia cohorts from different centers, totaling 96 and 13 patients respectively, were subsequently screened for *RFC1* resulting in a combined 705 subjects tested.

**Results:** In the initial cohort, 42 samples were identified with one expanded allele in the *RFC1* gene (7.0%), and 9 had two expanded alleles (1.5%). For the additional cohorts, we found 12 heterozygous samples (12.5%) and 7 biallelic samples (7.3%) in the larger cohort, and 1 heterozygous sample (7.7%) and 3 biallelic samples (23%) in the second. In total, 19 patients were identified with biallelic repeat expansions in *RFC1* (2.7%). Of these 19 patients, 6 (32%) had a clinical diagnosis of CANVAS, 10 had cerebellar ataxia with neuropathy (53%), and 3 had spinocerebellar ataxia (16%). No patients were identified with expansions in the *DAB1* gene.

**Conclusion:** In a large undiagnosed ataxia cohort from the United States, biallelic pathogenic repeat expansion in *RFC1* was observed in 2.7%. Testing should be strongly considered in ataxia patients, especially those with CANVAS or neuropathy.

## Introduction

Cerebellar ataxia is a heterogeneous disorder characterized by the inability to control gait and balance as well as coordinate voluntary muscle movement. Roughly 50% of affected individuals remain undiagnosed despite advanced genomic testing^1 2 3^. The most common genetic ataxias, as well as several rarer forms, are caused by nucleotide repeat expansions, which typically require non-sequence based testing to identify ^4 5, 6^. Current standard practice genetic evaluations for cerebellar ataxia involve testing for common repeat expansions followed by sequencing analysis for rare ataxias^2, 7^. The significant challenge to genetic testing for ataxic disorders involves the possibility of disease caused by rare or novel repeat expansion disorders that would be undetectable by such a strategy^3{Wallace, 2018 #3986^. Using non-parametric linkage analysis and whole genome sequencing, recent studies have identified a repeat expansion in the gene encoding Replication Factor C Subunit 1 (*RFC1*) related to certain forms of ataxia^8, 9^. This recessively inherited intronic (AAGGG) repeat expansion was shown to cause a specific ataxic phenotype referred to as Cerebellar Ataxia Neuropathy and Vestibular Areflexia Syndrome (CANVAS) ^8, 9^. In addition, this expansion has been suggested to be responsible for up to 22% (33/150) of cases of sporadic cerebellar ataxia and 63% (32/51) of cases of ataxia associated with sensory neuropathy^8^. To date, disease caused by the *RFC1* repeat expansion has been reported in Australia and United Kingdom but its incidence and prevalence in the United States remains undetermined.

Similarly, a dominant pathological repeat expansion was recently described in the intronic 5’ untranslated region of the DAB Adaptor Protein 1 (*DAB1*) gene causing spinocerebellar ataxia type 37 (SCA37) ^10^. In this disorder, a pentanucleotide (ATTTC) repeat insertion was identified within the normal (ATTTT) tandem repeat element. This *DAB1* repeat expansion was identified in patients from the southern Iberian Peninsula but remains unidentified elsewhere, including in the United States^10, 11^. The clinical heterogeneity of the dominant spinocerebellar ataxias are well-described^4^, making it unclear in what context SCA37 should be tested for in other regions of the world.

To address the frequency of these repeat expansion disorders in the United States, we assessed for the presence of expansions in *RFC1* and *DAB1* in a large cohort of 596 patients with unidentified cerebellar ataxia using fluorescent repeat primed polymerase chain reaction (RP-PCR). We identified biallelic *RFC1* expansion as causative for disease in 1.5% (n=9) of our patients and found no patients with a pathogenic *DAB1* expansion. We further tested two additional smaller cohorts from centers in different regions of the US and identified *RFC1-*mediated ataxia cases in 7.3% (7/96), and 23% (3/13) respectively for a total prevalence of 2.7% (19/705).

## Methods

### Patients

Patients were enrolled at the University of California, Los Angeles (UCLA) Ataxia Center, clinically assessed for acquired causes of ataxia, and then considered for genetic causes^2^. Only patients with negative testing for the common genetic ataxias (SCA1, SCA2, SCA3, SCA6, SCA7, and Friedreich Ataxia) were included in this study. All patients consented for DNA collection for genetic analysis. Peripheral blood was collected from patients and DNA was then isolated and purified using the Gentra Puregene Blood Kit (Qiagen) for genetic testing. The study methods used were approved by the UCLA Institutional Review Board. Additional deidentified DNA samples from 96 patients enrolled at the University of Chicago and 13 patients enrolled at the Brigham and Women’s Hospital under IRB approved procedures with identical inclusion criteria were subsequently tested as well.

### Repeat Expansion Testing

#### *RFC1* gene repeat expansion analysis

Fluorescent repeat-primed PCR (RP-PCR) was performed to detect *RFC1* pathogenic (AAGGG)_n_ alleles using previously published primers^8, 9^ with an optimized touchdown PCR protocol and Qiagen HotStarTaq. One primer set included the forward 5’FAM-ACTGACAGTGTTTTTGCCTGT-3’ primer, the anchor 5’-CAGGAAACAGCTATGACC-3’ primer, and the repeat 5’-CAGGAAACAGCTATGACCAAGGGAAGGGAAGGGAAGGGAAGGG-3’ primer that identifies the (AAGGG) repeats. A second primer set included the forward 5’FAM-TCAAGTGATACTCCAGCTACACCGT-3’ primer, the anchor 5’-CAGGAAACAGCTATGACC-3’ primer, and three repeat primers that identify the (AAGGG) repeats 5’-CAGGAAACAGCTATGACCAACAGAGCAAGACTCTGTTTCAAAAAAGGGAAGGGAAGGGAAGGGAA-3’, 5’-CAGGAAACAGCTATGACCAACAGAGCAAGACTCTGTTTCAAAAAGGGAAGGGAAGGGAAGGGAA-3’, and 5’-CAGGAAACAGCTATGACCAACAGAGCAAGACTCTGTTTCAAAAGGGAAGGGAAGGGAAGGGAA-3’. Fragment length analysis was performed using an Applied Biosystems 3730xl DNA Analyzer with Peak Scanner software (v. 2.0). Quantification and detection of non-pathogenic (AAAAG) motifs by RP-PCR was performed using the Takara LA PCR kit in combination with a touchdown PCR accommodating the recommended extension temperature for the enzyme. A fluorescently labeled forward primer (5’-ACTGACAGTGTTTTTGCCTGT) (10 μm), M13 anchor primer (5’-CAGGAAACAGCTATGACC) (10 μm) and (AAAAG) specific primer (5’CAGGAAACAGCTATGACC_AAAAGAAAAGAAAAGAAAAGAAAAG) (1 μm) were used to perform RP-PCR. Products were run on an ABI3730xl DNA analyzer and subsequently analyzed using GeneMarker v2.6.0 (SoftGenetics Inc.). To determine whether the genotype of samples with positive RP-PCR results were heterozygous or biallelic, standard PCR was performed using published primers^8^, forward 5’-TCAAGTGATACTCCAGCTACACCGTTGC-3’ primer and the reverse 5’ GTGGGAGACAGGCCAATCACTTCAG-3’ primer. Observation of a band at or near 348bp (wild-type size) corresponding to repeat sizes of less than approximately 60 repeats (approximately 650bp) identified patients as heterozygotes. As an internal control to prevent false positives, standard PCR was also performed simultaneously on the same sample with primers designed to amplify a 282 bp band from the *SPG11* gene (Supplemental Figure 1). All samples determined to have biallelic AAGGG expansions were assessed at least two times by both expansion RP-PCR testing and standard PCR genotyping in two separate laboratories.

**Figure 1.**
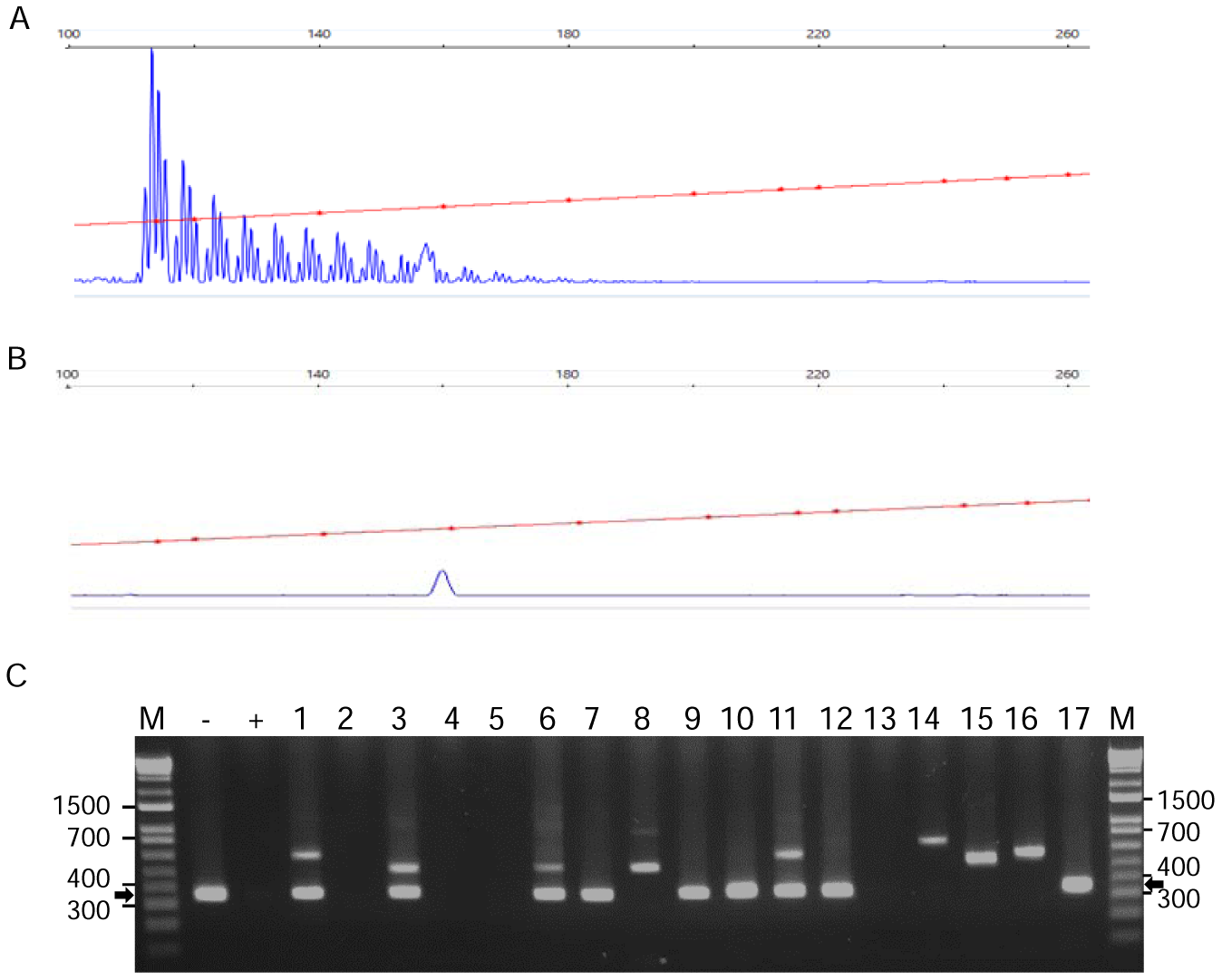
*RFC1* Expansion Analysis. Representative RP-PCR results from a patient with disease due to (A) biallelic expanded (AAGGG) pathogenic alleles or a control individual (B) with wild-type alleles. Samples with RP-PCR evidence of an expanded *RFC1* allele were genotyped by standard PCR for biallelic expansion (C). Standard PCR allows categorization of individuals as biallelic (no band, lanes 2, 4, 5, 13), heterozygous wild-type (348 bp band, arrow, lanes 7, 9, 10, 12, 17), or heterozygous with a non-pathogenic polymorphic expansion (variable sized band(s), lanes 1, 3, 6, 8, 11, 14, 15, 16). M, marker. -, wild type control. +, biallelic control.

#### *DAB1* gene repeat expansion analysis

Repeat-Primed PCR was performed to detect expansion of the normal (ATTTT) *DAB1* repeat region and for detection of the pathogenic (ATTTC)_n_ insertion using published primers^10 12^. Fragment length analysis was performed as described above.

### Data Availability

The datasets generated and/or analyzed during the current study are available from the corresponding author upon reasonable request.

## Results

We examined the prevalence of repeat expansions in *RFC1* and *DAB1* in a large cohort from the tertiary referral Ataxia Center at UCLA. The demographics of this 596 subject cohort are described in Table 1. Average age was 55 years, 50% of the patients were female, and 66% were white, non-Hispanic. The most common phenotypes were spinocerebellar ataxia (26.5%), pure cerebellar ataxia (21.6%), and multiple system atrophy (17.4%). To assess for the presence of pathogenic (AAGGG) repeat expansions in *RFC1*, fluorescent repeat-primed fragment analysis was performed and identified at least one expansion in 51 out of 596 patients (8.6%, Figure 1). Standard PCR was used to genotype subjects for the presence of a heterozygous or a pathogenic biallelic expansion. Nine subjects (1.5%) were found with biallelic expansions. Of these patients 3 subjects presented clinically with CANVAS (33%, 100% of phenotype), 5 had cerebellar ataxia with neuropathy (56%, 12% of phenotype), and 1 had spinocerebellar ataxia (11%, 0.6% of phenotype).

**Table 1.**
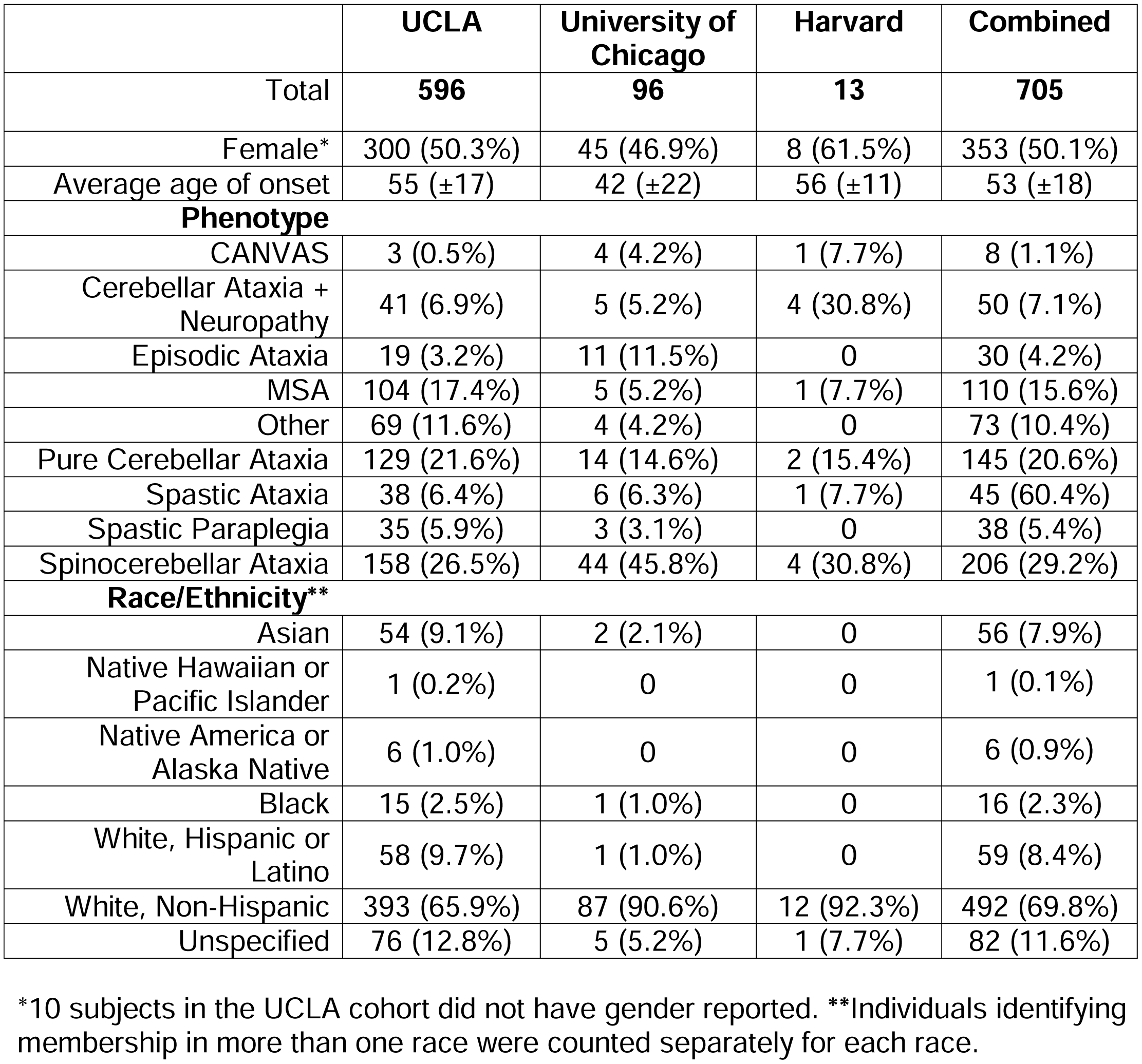
Patient Demographics

**Table 2.** Demographics of Patients with Biallelic Expansions in the *RFC1* Gene

| **Sample** | **Phenotype** | **Sex** | **Age of onset** | **Ethnicity** | **Inheritance** |
| --- | --- | --- | --- | --- | --- |
| **UCLA1** | CANVAS | Male | 78 | White, Non-Hispanic | Sporadic |
| **UCLA2** | Cerebellar Ataxia + Neuropathy | Female | 81 | White, Non-Hispanic | Familial |
| **UCLA3** | Cerebellar Ataxia + Neuropathy | Female | 66 | White, Hispanic or Latino | Familial |
| **UCLA4** | Cerebellar Ataxia + Neuropathy | Female | 71 | White, Non-Hispanic | Sporadic |
| **UCLA5** | Cerebellar Ataxia + Neuropathy | Male | 69 | White, Non-Hispanic | Sporadic |
| **UCLA6** | Cerebellar Ataxia + Neuropathy | Female | 71 | White, Non-Hispanic | Sporadic |
| **UCLA7** | CANVAS | Female | 69 | White, Non-Hispanic | Familial |
| **UCLA8** | CANVAS | Male | 63 | White, Non-Hispanic | Familial |
| **UCLA9** | Spinocerebellar Ataxia | Female | Unknown | Unspecified | Unknown |
| **UC1** | CANVAS | Female | 33 | White, Non-Hispanic | Familial |
| **UC2** | CANVAS | Male | 40 | White, Non-Hispanic | Sporadic |
| **UC3** | Cerebellar Ataxia + Neuropathy | Female | 42 | White, Non-Hispanic | Familial |
| **UC4** | Cerebellar Ataxia + Neuropathy | Female | 39 | White, Non-Hispanic | Familial |
| **UC5** | Spinocerebellar Ataxia | Male | 45 | White, Non-Hispanic | Sporadic |
| **UC6** | Spinocerebellar Ataxia | Female | 38 | White, Non-Hispanic | Sporadic |
| **UC7** | Cerebellar Ataxia + Neuropathy | Male | 38 | White, Non-Hispanic | Familial |
| **H1** | Cerebellar Ataxia + Neuropathy | Female | 55 | White Non-Hispanic | Sporadic |
| **H2** | CANVAS | Male | 60 | White Non-Hispanic | Sporadic |
| **H3** | Cerebellar Ataxia + Neuropathy | Female | 64 | White Non-Hispanic | Sporadic |

To validate these findings, we tested two additional smaller cohorts from centers in different regions of the US, the University of Chicago (UC) and Brigham and Women’s Hospital affiliated with Harvard Medical School. Demographics were similar to the UCLA cohort (Table 1). The larger UC cohort, consisting of both sporadic and familial cases, showed heterozygous expansions in 12 individuals (12.5%) and pathogenic biallelic expansion in 7 patients (7.3%). 2 subjects had CANVAS (29%, 50% of phenotype), 3 had cerebellar ataxia with neuropathy (43%, 60% of phenotype), and 2 had spinocerebellar ataxia (29%, 4.5% of phenotype). In the smaller Harvard cohort, 1 individual showed heterozygous expansion (7.7%) and 3 of 13 patients (23%) had pathogenic biallelic expansions. 2 of these had cerebellar ataxia with neuropathy (67%, 50% of phenotype) while 1 had CANVAS (33%, 100% of phenotype). Collectively, 19 of 705 subjects showed biallelic expansions across all cohorts for a total prevalence of 2.7% (19/705). All the patients were white (100%) and only 1 was Hispanic (5.3%). 10 of the cases showed sporadic onset (53%). In total, 6 subjects had CANVAS (32%, 75% of phenotype), 10 had cerebellar ataxia with neuropathy (53%, 20% of phenotype), and 3 had spinocerebellar ataxia (16%, 1.5% of phenotype).

For *DAB1* repeat expansion analysis, 83/596 (13.9%) subjects showed an expanded ATTTT allele by RP-PCR analysis however none of these patients possessed the pathogenic ATTTC insertion indicating that no patients within the cohort had SCA37.

## Discussion

Overall, in a large undiagnosed ataxia cohort of 705 patients from three tertiary referral centers in the United States, biallelic pathogenic (AAGGG) repeat expansions in *RFC1* were observed in 2.7% of patients. The observation that the majority of cases (95%) were white of European ancestry is consistent with previous reports^8, 9^. The high rate of heterozygosity (7.8%) in our cohort is notable but is similar to one previous study which calculated an allele frequency of 4% in control populations of 69 and 133 individuals based on haplotype in next-generation sequencing datasets ^9^, although another study found a frequency of 0.7% in a cohort of 304 healthy controls using RP-PCR^8^. Standard PCR analysis indicated that, under the conditions of our AAGGG RP-PCR assay, we were able to detect small expansions up to 60 repeats above wild-type (∼650bp, Figure 1, Supplemental Figure 1) and therefore we suspect that our high rate of detection may be due, in part, to the detection of confounding mildly expanded alleles below the estimated 400 repeat margin of pathogenicity ^8^ (Figure 1). It is interesting to note that small AAGGG expansions have not previously been reported in patients or controls^8, 9^, and since all subjects tested presented with some form of cerebellar ataxia, we cannot exclude the possibility that expanded AAGGG repeats may contribute to the development of cerebellar ataxia in some heterozygous individuals. We also cannot rule out a contribution of false positive detection of other small polymorphic non-pathogenic non-AAGGG repeats ^8^ in some heterozygous individuals. We did confirm all biallelic subjects with RP-PCR and standard PCR in at least two experiments each in two separate laboratories and further determined that none of these individuals harbored the most common non-pathogenic expanded repeat, AAAAG ^8^ (data not shown).

Additionally, the pathogenic (ATTTC) repeat in the *DAB1* gene, causative for SCA37, was not observed in our large undiagnosed ataxia cohort of 596 individuals of mostly white, non-Hispanic ancestry, consistent with the observation of a founder effect in the Iberian Peninsula^10, 11^ and suggesting that while this disorder should be considered in that population, it is likely extremely rare in the United States. Although no pathogenic ATTTC insertions were found, it is possible that extremely large repeats of ATTTT flanking a pathogenic ATTTC insertion might prevent amplification of products from the RP-PCR, so false negatives cannot be ruled out in this study, although this has not been commonly observed in published reports ^10 12^.

Consistent with prior reports, our study identified biallelic *RFC1* expansions in a high percentage of patients with CANVAS (n=6, 75%) and cerebellar ataxia with neuropathy^8^ (n=10, 20%). Although we do not have electrophysiological data on all subjects, of the biallelic patients identified, all appeared to have a large fiber neuropathy (data not shown), which would be an important focus for further clinical investigation. Additionally, we also observed biallelic expansion in a notable percentage of patients with generalized spinocerebellar ataxia (n=3, 1.5%), a frequency on par with the majority of rare ataxic disorders identifiable by clinical sequencing^2^. Taken together, these results suggest that *RFC1* expansion testing is high-yield in cases of CANVAS and cerebellar ataxia with neuropathy but should also be considered in the genetic workup of patients with undiagnosed spinocerebellar ataxia.

---

Supplemental Figure 1. Standard PCR Control.

To control for failure of the PCR reaction inadvertently resulting in a false positive result, standard PCR was performed simultaneously using *RFC1* primers (expected size of 348 bp for wild-type, black arrow) and control primers for *SPG11* (expected size of 282 bp, grey arrow). Lanes 1, 2, 3, 11, 13 represent biallelic expansions (no *RFC1* band). Lanes 5, 6, 14, 15, 16 represent wild type individuals. Lanes 4, 7, 8, 9, 10, 12, 17 represent heterozygous individuals with non-pathogenic polymorphic expansions. M, marker.

## Appendix 1 Author Contributions

| Name | Location | Role | Contribution |
| --- | --- | --- | --- |
| Dona Aboud Syriani | University of California, Los Angeles | Author | Acquisition of data, analysis of data, drafting manuscript. |
| Darice Wong, PhD | University of California, Los Angeles | Author | Analysis of data, study design, drafting manuscript. |
| Claudio M. De Gusmao, MD | Brigham and Women's Hospital and Harvard Medical School | Author | Acquisition of data, revising manuscript for intellectual content. |
| Sameer Andani BS | University of Chicago, Chicago | Author | Acquisition of data, revising manuscript for intellectual content. |
| Yuanming Mao BS | University of California, Los Angeles | Author | Acquisition of data, analysis of data, drafting manuscript for intellectual content. |
| Giacomo Glotzer | University of Chicago, Chicago | Author | Acquisition of data, revising manuscript for intellectual content. |
| Paul J. Lockhart, PhD | Murdoch Children's Research Institute | Author | Study design, revising manuscript for intellectual content. |
| Sharon Hassin-Baer, MD | Sackler Faculty of Medicine, Tel-Aviv University, Tel-Aviv, Israel | Author | Acquisition of data, revising manuscript for intellectual content. |
| Vikram Khurana, MD, PhD | Brigham and Women's Hospital and Harvard Medical School | Author | Acquisition of data, study design, revising manuscript for intellectual content. |
| Soma Das, PhD | University of Chicago, Chicago | Author | Data analysis, study design, revising manuscript for intellectual content. |
| Christopher M. Gomez, MD, PhD | University of Chicago, Chicago | Author | Acquisition of data, study design, analysis of data,<br>revising manuscript<br>for intellectual<br>content. |
| Susan Perlman,<br>MD | University of California,<br>Los Angeles | Author | Acquisition of data,<br>revising manuscript<br>for intellectual<br>content. |
| Brent L. Fogel, MD,<br>PhD | University of California,<br>Los Angeles | Author | Study design and<br>supervision,<br>acquisition of data,<br>analysis of data,<br>drafting and<br>revising manuscript<br>for intellectual<br>content. |

## Acknowledgements

The authors wish to thank all the patients and their families for their participation in this study.

## Notes

Study Funding: Funding: This work was supported by the National Institute for Neurological Disorders and Stroke [R01NS082094 to BLF]. BLF acknowledges support through donations to the University of California by the Rochester Ataxia Foundation. PJL was supported by the Vincent Chiodo Foundation. CMG and VK acknowledge support from the National Ataxia Foundation and the Brigham Research Institute.

