## Supplementary figures and images for "Prevalence of *RFC1*-Mediated Spinocerebellar Ataxia in a United States Ataxia Cohort"

### Supplemental Figure 1

#### Slide 1
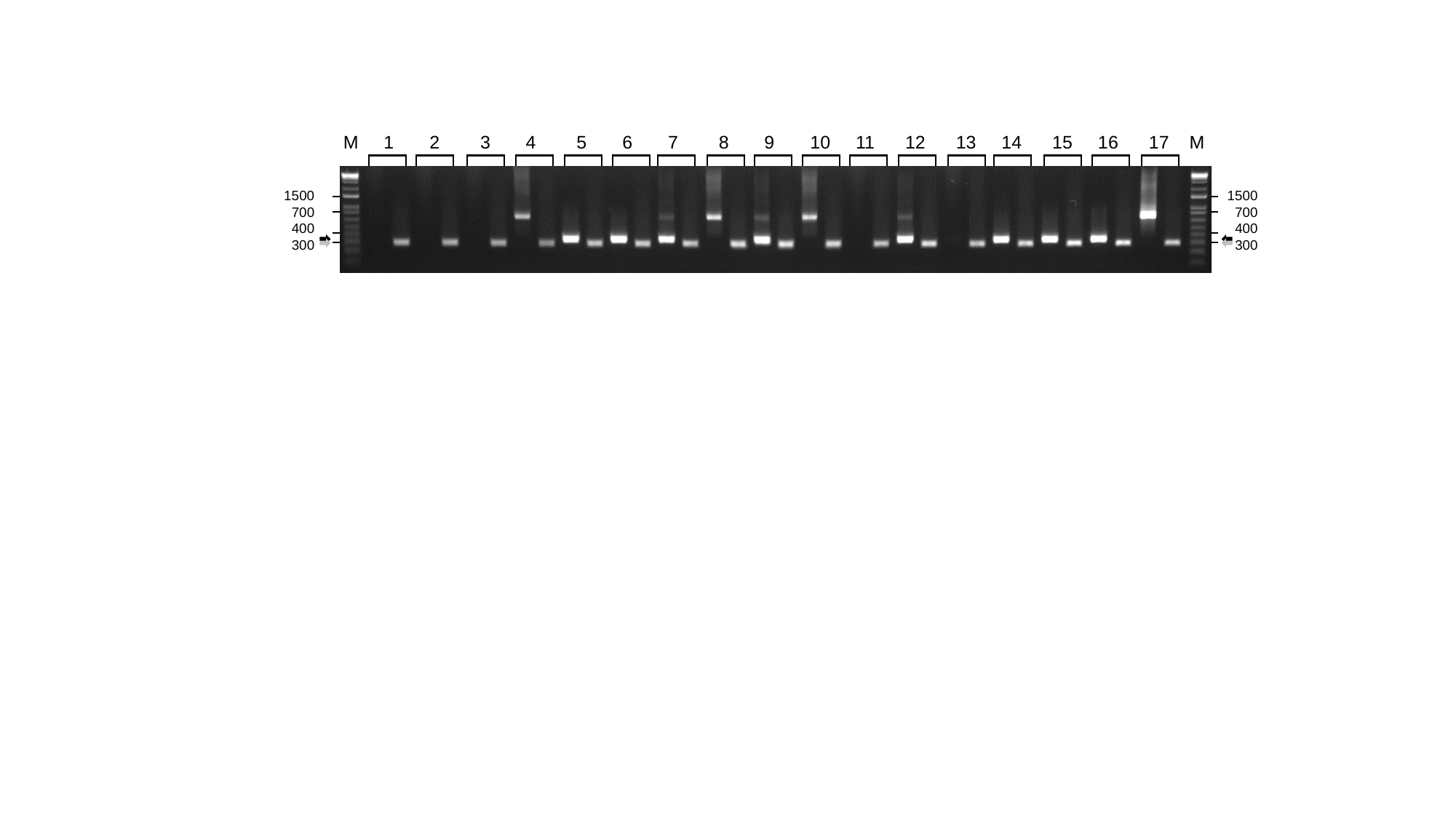

M 1 2 3 4 5 6 7 8 9 10 11 12 13 14 15 16 17 M
1500
700
400
300
1500
700
400
300
